# Bounds on the computational complexity of neurons due to dendritic morphology

**DOI:** 10.1101/2025.07.11.664448

**Authors:** Anamika Agrawal, Michael A. Buice

## Abstract

The simple linear threshold units used in many artificial neural networks have a limited computational capacity. Famously, a single unit cannot handle non-linearly separable problems like XOR. In contrast, real neurons exhibit complex morphologies as well as active dendritic integration, suggesting that their computational capacities outperform those of simple linear units. Considering specific families of Boolean functions, we empirically examine the computational limits of single units that incorporate more complex dendritic structures. For random Boolean functions, we show that there is a phase transition in learnability as a function of the input dimension, with most random functions below a certain critical dimension being learnable and those above not. This critical dimension is best predicted by the overall size of the dendritic arbor. This demonstrates that real neurons have a far higher computational complexity than is usually considered in neural models, whether in machine learning or computational neuroscience. Furthermore, using architectures that are, respectively, more “apical” or “basal” we show that there are non-trivially disjoint sets of learnable functions by each type of neuron. Importantly, these two types of architectures differ in the robustness and generality of the computations they can perform. The basal-like architecture shows a higher probability of function realization, while the apical-like architecture shows an advantage with fast retraining for different functions. Given the cell-type specificity of morphological characteristics, these results suggest both that different components of the dendritic arbor as well as distinct cell types may have distinct computational roles. Our analysis offers new directions for neuron-level inductive biases in NeuroAI models using scalable models for neuronal cell-type specific computation.

## 1 Introduction

The relationship between neuronal structure and function has received renewed attention due to the growing appreciation of morphoelectric diversity across cell types [Gouwens et al., 2019, 2020, Dembrow et al., 2024, Kalmbach et al., 2021]. Yet, other than classical studies of synaptic integration and circuit motifs, the computational roles of single neurons, shaped by their dendritic morphology, remain underexplored. While traditional models often simplify neurons as linear threshold units, such approximations neglect the rich, nonlinear processing capabilities arising from the spatial and electrical organization of dendritic arbors. Recent experimental evidence demonstrates that dendrites actively integrate inputs in a manner that is both cell-type specific and nonlinear [London Preprint. and Häusser, 2005]. These findings suggest that dendritic and morphological properties endow neurons with complex computational capabilities, positioning them between simple linear classifiers and deep artificial networks [Cazé et al., 2013, Bicknell and Häusser, 2021].

It has been shown that single neurons can transcend passive linear integration and compute nonlinearly separable functions, such as the XOR function [Gidon et al., 2020] and feature-binding tasks [Bicknell and Häusser, 2021]. However, these studies typically focus on specific nonlinearly separable problems and therefore do not quantify the broader computational limits of single neurons across a range of function complexities. Prior work [Cazé et al., 2013] demonstrated that neurons with local nonlinearities can leverage different integration strategies to solve certain nonlinearly separable tasks. Building on this foundation, we employ Boolean functions to systematically scale the complexity of computational problems with the dimension of input. In addition, we explore specific families of Boolean functions to characterize the capabilities of the dendritic arbor.

In this work, we examine how dendritic morphology constrains the upper bounds of computational complexity achievable by a single neuron. We model neurons as abstract networks of non-linearly integrating dendritic compartments, organized according to archetypal branching architectures. Within this framework, dendritic networks are tasked with learning Boolean functions, allowing precise control over input complexity and objective difficulty. By assessing function realizability across different input dimensions and measures of Boolean function complexity (here, entropy [Mingard et al., 2019] and sensitivity [Kenyon and Kutin, 2004]), we systematically relate morphological features, such as breadth and depth of dendritic trees, to the space of computations that single neurons can perform.

To ground this analysis, we compare two canonical morphologies: (1) a shallow, broad topology reminiscent of basket cells and basal dendritic tufts [Häusser and Mel, 2003, Tzilivaki et al., 2019], and (2) a deep, narrow topology modeled after the apical tufts of excitatory neurons [Häusser and Mel, 2003, Spruston, 2008]. Despite containing the same number of nonlinearly integrating compartments, these architectures exhibit distinct computational strengths: shallow networks excel at learning low-dimensional, low-sensitivity functions, while deep networks generalize more robustly and retrain more efficiently across complex tasks. Importantly, we show that dendritic architectures impose a critical input dimensionality threshold, beyond which the probability of learning a randomly sampled Boolean function collapses sharply. This behavior is reminiscent of phase transitions in circuit complexity (e.g., phase transition in 3-SAT, [Karp, 2009]) and highlights the intrinsic computational limits imposed by morphology. Furthermore, different dendritic architectures preferentially realize distinct subsets of Boolean functions, suggesting a possible division of computational labor across cell types within neuronal ensembles. By abstracting dendritic trees into structured, deep network-like architectures, our work offers a biologically grounded and tractable framework for understanding how single-neuron computations are shapedand limitedby morphology. Incorporating such cell-type-specific computations into models of neural circuits opens new avenues for rethinking both theoretical and systems neuroscience paradigms.

## 2 Testing the limits of single neuron computation with abstract models of dendritic morphology

**Abstract Model of a Nonlinear Dendritic Network** We model a single neuron as a feedforward network comprising individual dendritic rectification units, which are arranged according to the morphology of the dendritic arbor of the neuron. (Figure 1 a). Every independent dendritic compartment *b* is modeled as a unit in an equivalent multi-layer network representation of the neuron. The distal branches sum and rectify the n-dimensional input layer (*x*_1_, *x*_2_, *x*_*n*_) as:

**Figure 1.**
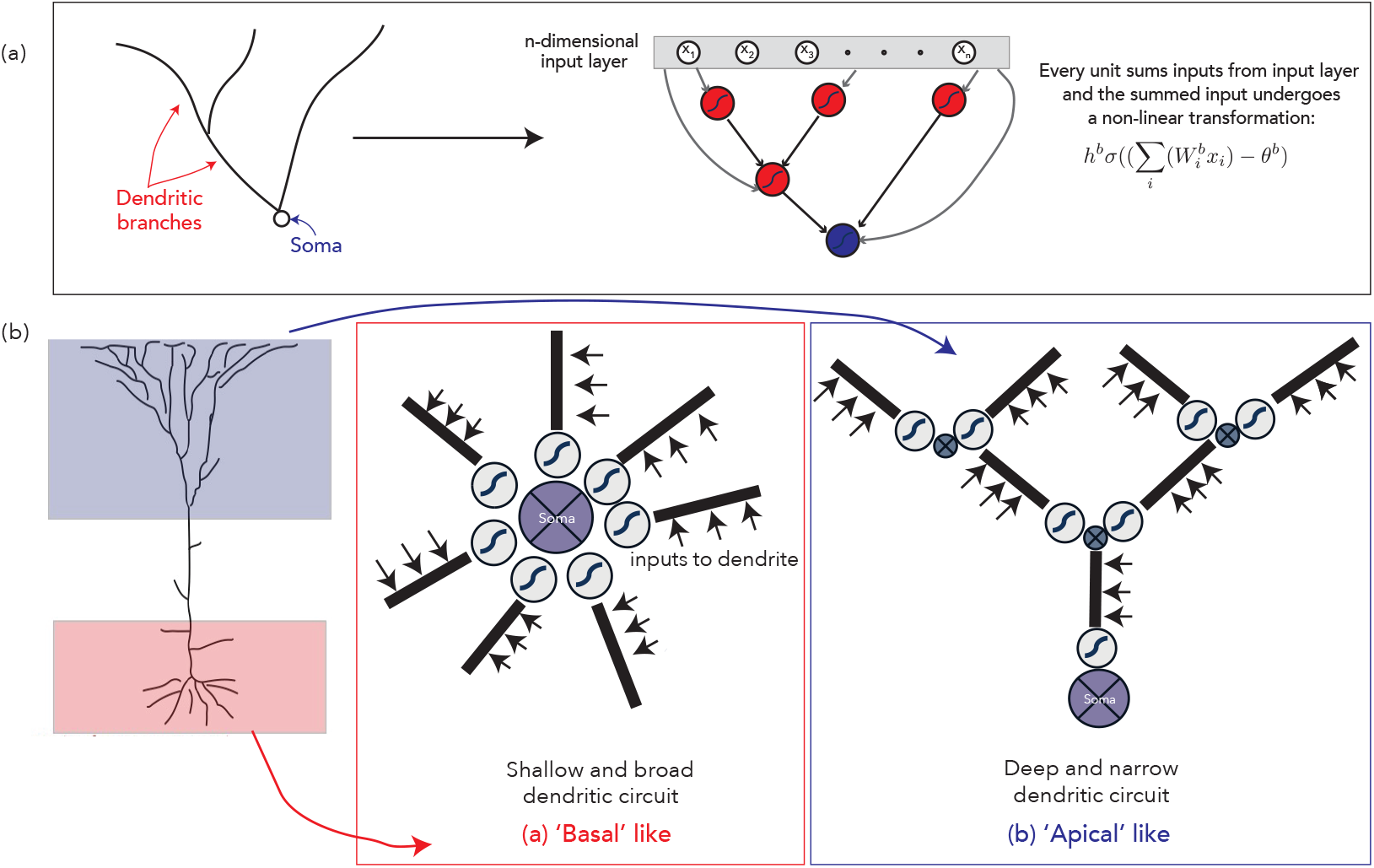
(a) Abstraction of dendritic morphology. (b) Two dendritic architectures: (1) Shallow and broad: all nonlinear compartments connect directly to the soma, resembling basal tufts and enabling parallel processing. (2) Deep and narrow: nonlinear compartments are arranged hierarchically, resembling apical tufts with electrotonically distinct subtrees.

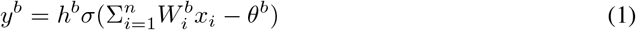

Where *y*^*b*^ is the output of the dendritic branch *b, h*^*b*^ and *θ*^*b*^ are the branch-specific gain and threshold parameters, 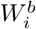 is the weight matrix element for input *i* on branch *b*, and *σ* is the logistic sigmoid function.

If a branch *b*′ also receives inputs from branches *b* with in-degree connections to it as well as from direct inputs, its output is given by:

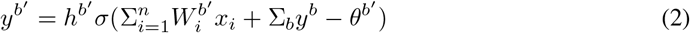

The soma is assumed to be another node which rectifies its inputs from attached dendrites and the input layer:

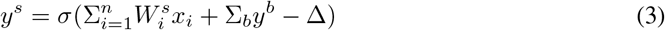

where Δ is the soma threshold parameter. Schematics of these models are presented in Figure 1 a, with the input layer forming ‘skip connections’ to represent that all nodes of the network receive input from the input layer.

For our first set of analyses, we investigate how differences in dendritic arrangement impact the local computational capabilities of single neurons. For simplicity, we used two extremes of dendritic topology arrangements. These architectures reflect trade-offs in breadth and depth of a dendritic circuit especially when there are biophysical limits on the extent of the dendritic arbor. Most neuronal cell types consist of these two dendritic tufts with different levels of dendritic branch units dedicated to each tuft.

The first model is one with a single hidden layer that represents multiple dendritic compartments directly connected to the soma, constituting a broad but shallow architecture. This arrangement is reminiscent of basket cells [Tzilivaki et al., 2019] and the basal tuft of certain classes of neurons (Spruston [2008], Figure 1B). This parallel arrangement is henceforth referred to as the ‘basal / parallel’ type. The second extreme focuses on architectures with additional hidden layers, denoting the hierarchical nature of tree-like connections in some dendritic tufts. Such an equivalent architecture, if constrained by the number of available dendritic compartments, will be deeper but narrower than the basal model. This arrangement is reminiscent of the apical tuft of certain excitatory cell types (Spruston [2008], Figure 1B), and is henceforth referred to as the ‘apical’ type. We will typically compare these models with the same number of compartments.

For simplicity, we first consider a very small dendritic circuit with *N*_*branch*_ = 7 in both architectures. (Figure 1 B). We compare both of these dendritic models to a control linear integration model, that merely assumes that all branches perform linear summation on the inputs with a final non-linear rectification step at the soma:

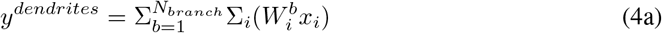

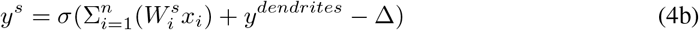

### Characterizing the Computational Complexity of a Dendritic Networ

The computational complexity of a Boolean function, given a fixed set of gates such as AND and OR, can be characterized by the number of gates as well as the minimum depth in a circuit needed to implement the function [Wegener, 1987]. Here, we ask, given a particular multi-layer network, and a particular family of Boolean functions: what are the limitations on the network for implementing that family of functions?

A single neuron can be thought of as receiving inputs from *N*_*dim*_ input streams that could represent different sources of information. If the input *X*_*i*_ is 0, that means that the input stream is inactive with low, noisy firing, and if it is 1, it can be interpreted to mean that the input stream is active and has a sufficiently high signal-to-noise ratio. The 0/1 inputs could also be interpreted as very low/finite EPSPs from distinct input sources to the neuron.

Previous work on dendritic learning with increasing size of input stream has focused only on memory capacity [Poirazi and Mel, 2001], i.e., the number of patterns in a space of all possible patterns in a given input dimension *N*_*dim*_ that can be learned by a single neuron. In our analysis, we instead test the capacity of a dendritic circuit to make arbitrary classifications, specified by the truth table of a boolean function. It has been famously shown [Gardner and Derrida, 1988] that the number of patterns in a given input stream dimension *N*_*dim*_ that can be learned by a linear perceptron (i.e. the fraction of linearly separable problems) scales as *p* = *a* ∗ *N*_*dim*_, *a <* 1. Therefore, as the size of the input stream *n* increases, the fraction of total problems that are linearly separable 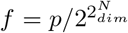 decreases to an extent that the probability that a linear perceptron can solve a randomly sampled function in this space becomes close to 0. This is consistent with the fact that the computational complexity of a function also increases as *n* increases.

We consider Boolean functions of sufficiently high dimension that it would prohibitive to test them all. Therefore, we consider multiple families of functions defined by constraints on the functions in question. First, we consider the class of ‘typical’ functions in *N*_*dim*_, the functions randomly sampled by choosing the output independently and identically for each input pattern, with probability *p* = 0.5 of returning 1 and likewise for 0. These typical Boolean functions tend to have high sensitivity (defined in the subsequent paragraphs). For a given input dimensionality *N*_*dim*_, there are 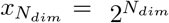 possible input combinations, and 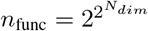 Boolean functions. In the face of extremely large number of possible functions in input dimensions higher than 4, the class of typical functions is to sample the space of “common” functions. We test the performance of the three model dendritic architectures, (1) basal-like (2) apical-like and (3) linear-like on this class of functions over a range of *N*_*dim*_.

Second, we consider the class of Boolean functions of fixed entropy, defined simply as the fraction min(number of input rows that map to 1, number of input rows that map to 0)/total number of input rows in the truth table [Mingard et al., 2019]. These functions are generated randomly by considering values of *p*! = 0.5.

Finally, we consider the class of Boolean functions of fixed sensitivity [Rubinstein, 1995, Mossel et al., 2005]. The sensitivity of a Boolean function in dimension *n* is defined as follows:

#### Definition 2.1

(Def 1). Let *f* : 0, 1^*n*^ → 0, 1 be a Boolean function and let *x* ∈ 0, 1^*n*^. The sensitivity of *f* at *x* is the number of positions *i* ∈ [*n*] such that flipping the *i*^*th*^ bit of *x* changes the output *f* (*x*). The sensitivity *s*(*f*) of *f* is the maximum sensitivity *s*(*f, x*) over all points *x*.

Informally, the sensitivity of a function *f* measures how unstable the output of *f* is to small perturbations to the input. We observe that both entropy and sensitivity increase monotonically with respect to each other (Supplemental Figure 5) and also scale the difficulty in learnability for the linear dendritic architecture (Figure 3 A). A low sensitivity or low entropy Boolean function can be thought of as a function with some non-influential bits that do not impact the truth table [Kalai, 2016]. These functions could be analogous to neural computations with high-dimensional input but with various nuisance or irrelevant features. Alternatively stated, they are effectively low dimensional functions compared to the input. Since the sensitivity of a Boolean function increases monotonically with its entropy, we used the same set of functions with varying entropy and calculated their sensitivity. We independently verified that generating Boolean functions of fixed sensitivity via rejection sampling gives similar distributions of functions.

After defining these classes of Boolean functions, we test whether or not each class of functions can be learned by a given architecture. A function mapping is considered learnable by a given architecture, if a set of weights 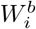 and non-linearities *θ*^*b*^, *h*^*b*^ can be learned by that architecture during training to map the input 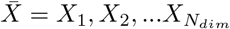 to its boolean output 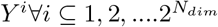, where *i* denotes the index of the row of the truth table representing the boolean function. To quantify this, we use the Hamming distance between the thresholded and binarized output of the dendritic network after training on the truth table Supplemental Figure 1 details our training approach.

For a given class of functions, suppose there are *N*_*func*_ functions of the class that the architecture is trained on over *n*_*trials*_ trials with different initial conditions. We utilize three different learning metrics to test the learnability of the class:

1. The mean Hamming distance between the network output and training target over trials,

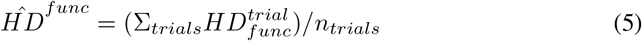

The mean and variance of this quantity over the sampled functions *N*_*func*_ is then visualized.
2. Function learning probability,

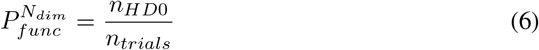

is the probability of learning a particular random function and *n*_*HD*0_ is the number of trials where the given function was learned with a hamming distance ~ 0 in *n*_*HD*0_. This is the fraction of times the network learned the function perfectly.
3. The ‘best-case’ learning outcome, i.e. the probability of typical function learning is instead calculated as

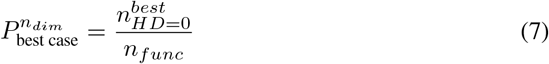

where 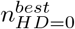 is the number of functions with a hamming distance of 0 in at least one of the trials tested. This is the fraction of functions for which there is at least one successful trial.

## 3 A phase transition in functional realizability

First, we trained the 3 dendritic networks (basal, apical, linear) on Boolean functions with input dimensionality varying from *N*_*dim*_ = 3 to *N*_*dim*_ = 8. Convergence was ensured by tuning the learning rate and the number of epochs for a given *N*_*dim*_. The training time for a given architecture and function over one set of initial conditions (trial) took a maximum of ~ 4 hrs (*N*_*dim*_ = 8). For multiple trials and functions, we use parallel computing over a cluster to run training on multiple functions and trials simultaneously.

For the first test, we randomly sample up to *n*_*func*_ = 1000 in a given input dimension. As anticipated, a linear dendritic circuit with only a somatic non-linearity fails to solve most functions even in the lower tested input dimension of *N*_*dim*_ = 3 (Supplemental Figure 2), while the basal and apical type learn most of the functions in *N*_*dim*_ = 3. To mitigate the effects of single-trial testing, we sampled a limited number of randomly-sampled typical functions (*n*_*func*_ = 100) across multiple trials (*n*_*trials*_ = 10), with each trial randomizing the initial conditions of dendritic parameters and weights. Figure 2 A shows the trends in the mean hamming distance across trials as *N*_*dim*_ increases. The mean hamming distance was computed across trials, as explained in 5. As anticipated, the mean hamming distance across trials increases as *N*_*dim*_ increases. However, the linear type has a significantly higher mean hamming distance across dimensions.

**Figure 2.**
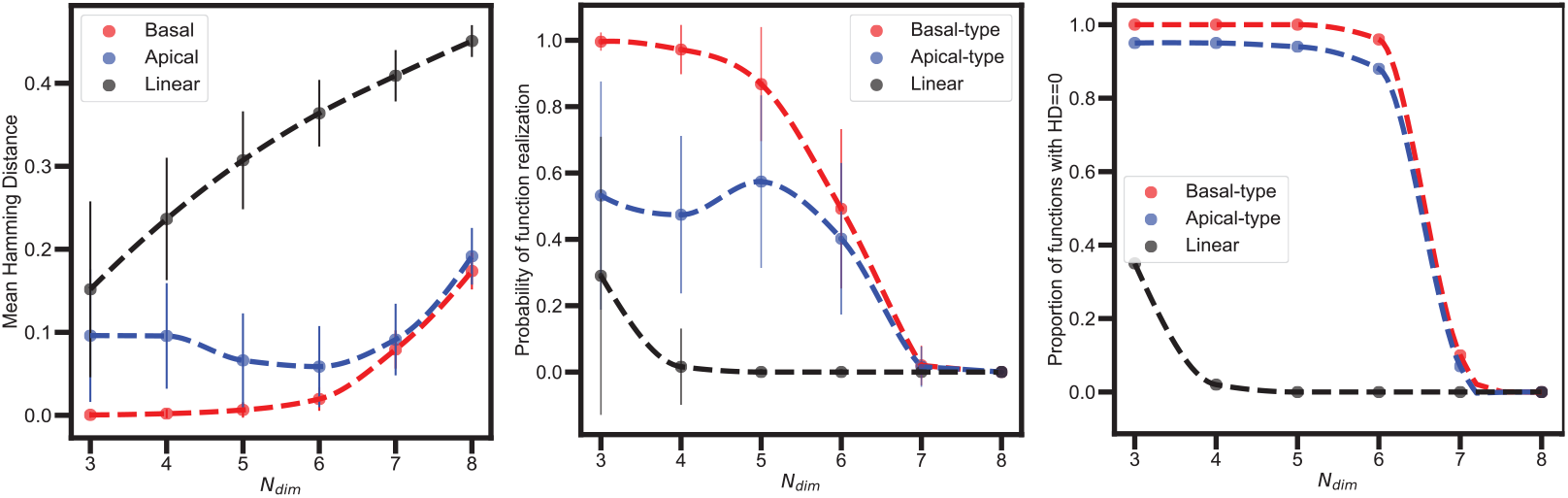
Phase transition in function realizability as *N*_*dim*_ increases. (a) Mean Hamming distance across *n*_*trials*_ = 10 for *N*_*func*_ = 100, calculated as in 5. Probability of function realization (6). (c) Best-case realizability (7). Apical and basal architectures show transitions at higher *N*_*dim*_ than the linear model.

In Figure 2 B, we visualize the probability of function realizability for the typical function class (Equation 7). Here, we observe an interesting transition from high probabiltiy of realizability to very low realizability probability for all architectures. This transition is even more evident in Figure 2 C, where the best-case learning outcome (Equation 7) is considered. While the linear architecture demonstrates this transition in a low input dimension (*N*_*dim*_ = 4), we observe this transition for basal and apical types as well for *N*_*dim*_ = 7.

It is known that given enough number of non-linearly integrating compartments (*N*_*branch*_ in the dendritic architecture shown in 1(a)), it is possible to solve any complexity of function [Cybenko, 1989]. However, in the case where we have a limited number of such compartments, we observe that there is a steep transition in the probability of realizability of a typical function in *N*_*dim*_ for the ‘basal’ architecture. We also verified that this transition is not dependent on the optimization mechanism (backpropagation in our case) by also testing DIRECT, [Jones et al., 1993, Gablonsky and Kelley, 2001], a non-backprop mechanism that does a grid-search style optimization (Supplemental Figure 4). Beyond the observed transition point, problems become so intractable that the probability of such a non-linear circuit solving it is close to 0.

In Supplementary Figure 3, we analyze how increasing dendritic compartments affects computational capacity. For the basal architecture, size increases by adding branches; for apical, we sample 2-layer tree-like circuits with increasing nodes. In both cases, capacity scales exponentially with compartment number. Because connections are feedforward and sparse, deeper networks trade off breadth, explaining why apical and basal architectures show similar size-cost trade-offs for increasing input dimensionality.

The apical architecture shows lower typical function realizability in low dimensions and greater variability across functions (Figure2B), though it matches basal performance in higher dimensions. This may stem from sensitivity to initial conditions, as uniform weight initialization may disadvantage hierarchical structures. The best-case analysis (Figure2C) compensates for this, revealing similar overall trends. We examine this variance further in the next sections.

For some Boolean functions, mismatches in output may be tolerable - e.g., if certain inputs are irrelevant or later corrected by additional signals. Given the high dimensionality of real dendritic inputs, partial learning may suffice. To test this, we relaxed the learning criterion to allow limited output mismatches (Supplementary Figure4) and found that in higher dimensions, the apical architecture shows slightly better approximate learning than the basal type.

## 4 Morphology based limits on single neuron computation

As observed in the error bars in Figure 2 B, the sampled functions in a given dimension can still have differing characteristics that can lead to variation in the probability of realizability within a given input dimension *N*_*dim*_. This variation is quite visible even within the *n*_*func*_ = 100 functions sampled in Figure 2. This required another order parameter that can differentiate boolean functions in the same *N*_*dim*_ in terms of their realizability difficulty by neural networks.

We go beyond the typically sampled functions to broaden the space of tested functions, by varying the entropy and thus the sensitivity of boolean functions sampled in a given input dimension, whose definitions and intuitive meaning are elaborated earlier. Supplemental figure 6 demonstrates how the distribution of sensitivity changes from 100 typical functions to 1000 functions with distributed values of sensitivity, showing that the entropy-based sampling scheme successfully samples functions with a varying degree of sensitivity.

Figure3B shows learning probabilities for 1000 sampled functions plotted by sensitivity, comparing basal (x-axis) and apical (y-axis) architectures. The two cell types specialize in different function classes: functions above the diagonal are better learned by apical cells, while those below are better learned by basal cells. In Figure3C, binning by sensitivity reveals that basal cells excel at learning low-sensitivity functions, but their performance drops as function complexity increases. In contrast, apical cells show a more uniform (but generally lower) learning probability across sensitivity levels, suggesting they imperfectly learn simple functions under standard training conditions. This pattern implies that apical cells generalize broadly but less precisely, whereas basal cells learn robustly with minimal generalization. We hypothesize that the apical architecture prioritizes generalization before fine-tuning, while the basal architecture learns functions independently and more deterministically. We test this idea further in the next section.

**Figure 3.**
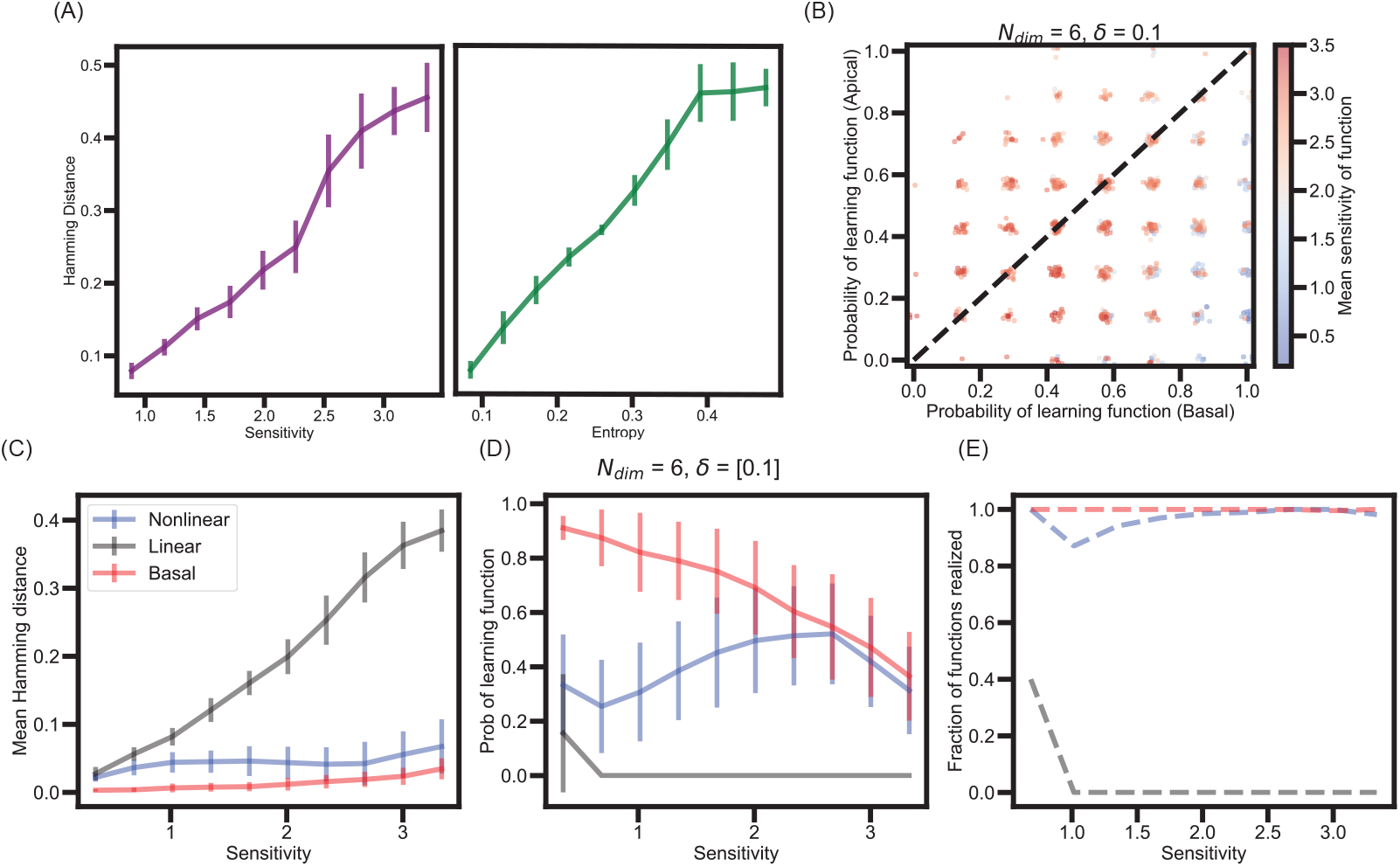
(A) Hamming distance increases with sensitivity and entropy for linearly integrating neurons, indicating greater difficulty in realizing such functions. (B) Probability of learning for 1000 functions (at *N*_*dim*_ = 6), plotted for basal (x-axis) vs. apical (y-axis) architectures. Points above or below the diagonal indicate architecture-specific preference. (CE) Mean Hamming distance, learning probability, and best-case realizability binned by function sensitivity, averaged over *n*_*trials*_ = 10.

## 5 Morphology based trade off between computational robustness and flexibility

As noted earlier, the apical architecture shows lower realizability under random initial conditions, particularly for low-sensitivity functions. To test whether this is due to different learning strategies, we assessed retraining flexibility (Figure 4). Both apical and basal architectures were first trained on *n*_func_ = 50 Boolean functions at *N*_dim_ = 5, with entropy increasing by 0.1 across batches of 10. We selected this dimensionality just below the realizability threshold to ensure all functions were learnable within a few trials. For each function, we saved the weights from a successful trial (*HD* = 0).

**Figure 4.**
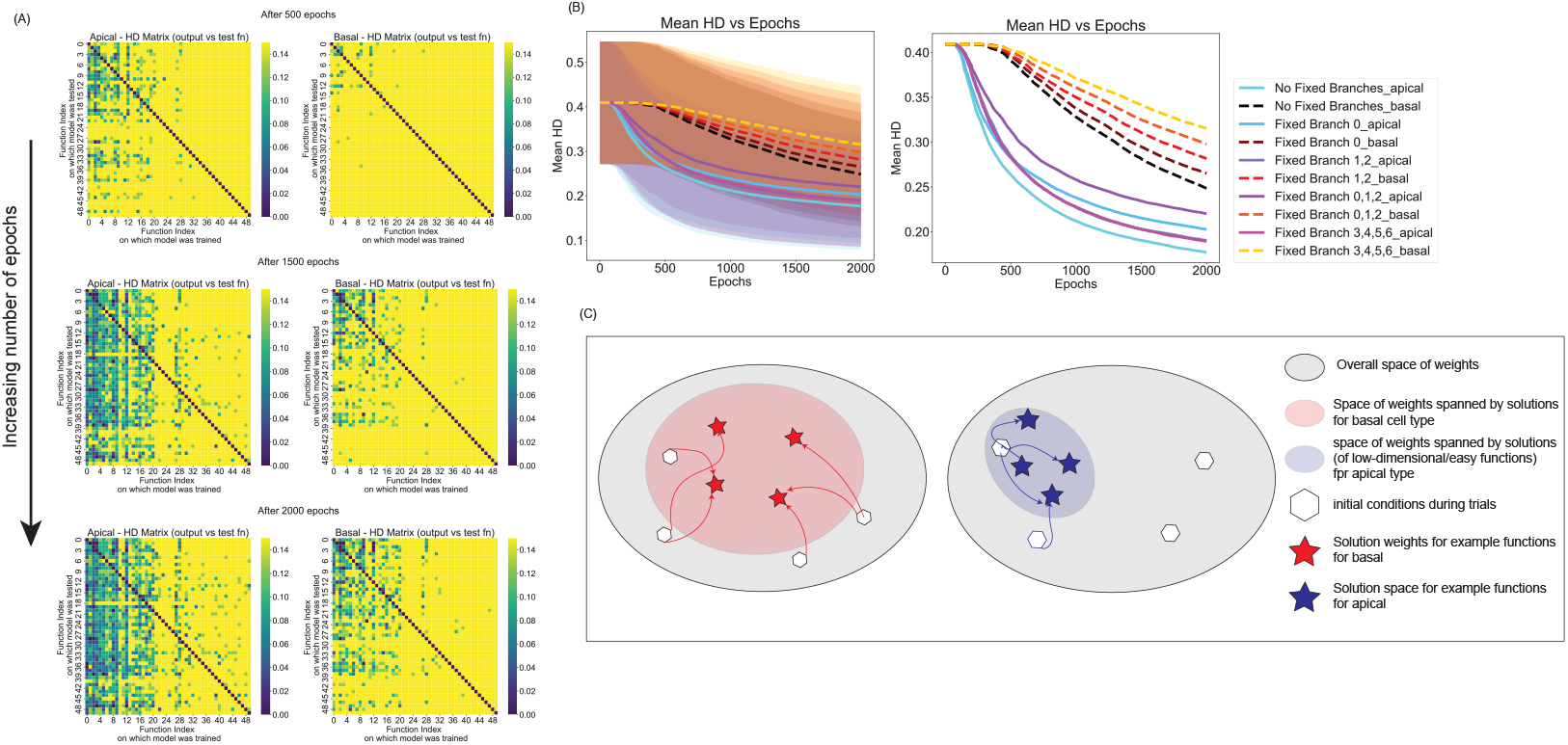
Apical architecture enables more flexible retraining. (A) Hamming distance *HD*_*j*→ *i*_(*t*) between pre-trained function *j* and test function *i* over increasing epochs (*t* = 500, 1500, 2000) shows faster convergence for apical cells, especially when pre-trained on low-entropy functions (*j* = 1–10). (B) Mean Hamming distance over epochs under different weight-freezing conditions. Apical performance is less affected when distal branches (36) are fixed, indicating their role in early generalization. (C) Schematic: basal cells are more robust across initializations; apical cells more flexible once pre-trained on simple functions.

Next, each architecture was retrained on function *i* using initial weights from function *j*, and we tracked Hamming distance *HD*_*j* → *i*_(*t*) over epochs *t*. As shown in Figure 4A, the diagonal elements (*j* = *i*) are zero by construction, but pretraining on low-entropy functions (*j* = 1–10) also accelerates learning for other functions. This effect is stronger for the apical architecture, which converges faster than basal across all *j* (Figure 4B).

We further tested retraining while fixing specific branch weights. Holding distal branches (*b* = 3–6) fixed had minimal impact on apical retraining, but significantly slowed basal retraining, where all branches are equivalent. Additional tests fixing both weights and nonlinearities (*h*_*b*_, *θ*_*b*_) of distal branches confirmed that only proximal branches need tuning for the apical cell to re-learn (Supplemental Figure 7).

These findings suggest that apical cells generalize using distal branches trained on simple functions (e.g., AND/OR), which serve as reusable building blocks for more complex tasks. In contrast, basal cells lack this generalization and must relearn each function independently. Thus, apical architectures are more flexible but less robust, while basal architectures are robust but less adaptable.

## 6 Discussion

Our results demonstrate that dendritic morphology is not merely a structural feature but a fundamental determinant of a neuron’s computational capacity. By abstracting dendritic trees into structured, resource-limited deep network architectures, we show that the arrangement and number of nonlinear integrating compartments impose strict bounds on the input-output functions a single neuron can realize. While nonlinear dendritic integration greatly expands computational capabilities beyond traditional linear threshold models, this expansion is not unbounded. We identify a sharp transition in functional realizability as input dimensionality increases: beyond a critical dimension, the probability of learning random Boolean functions collapses. This phase transition mirrors phenomena in circuit complexity theory, such as the 3-SAT satisfiability transition, revealing inherent computational limits shaped by morphological constraints.

Importantly, dendritic architecture mediates different computational trade-offs. “Basal-like” shallow, broad architectures favor robust, condition-insensitive computation, reliably learning simple functions across diverse initializations. In contrast, “apical-like” deep, narrow architectures favor flexibility: though initially less robust, once properly initialized, they retrain and generalize across a broader range of functions with fewer adaptations. These complementary properties suggest a natural division of computational labor across cell types: robust processors for stable operations and flexible processors for dynamic reconfiguration.

These findings have immediate implications for understanding cortical circuit organization. Excitatory neurons with elaborate apical tufts, such as layer 5 pyramidal neurons, may leverage their architecture to flexibly adapt to changing cognitive demands, while inhibitory neurons with simpler dendritic arbors ensure stability and reliability. Modeling efforts incorporating these single-cell morphological constraints could yield more accurate predictions of learning, plasticity, and resilience in large-scale cortical networks.

Our framework simplifies many aspects of neuronal computation. We focus on feedforward architectures with rectified nonlinearities, abstract away spiking dynamics, recurrent connections, and inhibition, and use Boolean function learning as a proxy for computational complexity. These abstractions omit key biophysical phenomena such as voltage-gated dynamics, synaptic plasticity, and temporally structured activity patterns. Nevertheless, the qualitative insights remain robust: morphology directly shapes the computational strategies available to a neuron.

By linking dendritic morphology to single-neuron computational limits via deep network architectures, we provide a tractable, scalable approach bridging cellular neuroscience with theoretical frameworks from machine learning and computational complexity. Future work extending this to ensembles of diverse neurons could reveal how local computational trade-offs influence circuit-level dynamics, shaping cognition, learning, and dysfunction. In short, morphology sculpts computation, shaping not only what neurons do, but how they learn and adapt. Table 1 summarizes how dendritic architectures impose distinct trade-offs between robustness, flexibility, and sensitivity. Morphological constraints shape the computational landscape, suggesting evolution has optimized not just wiring efficiency, but computational strategy at the cellular level.

**Table 1:**
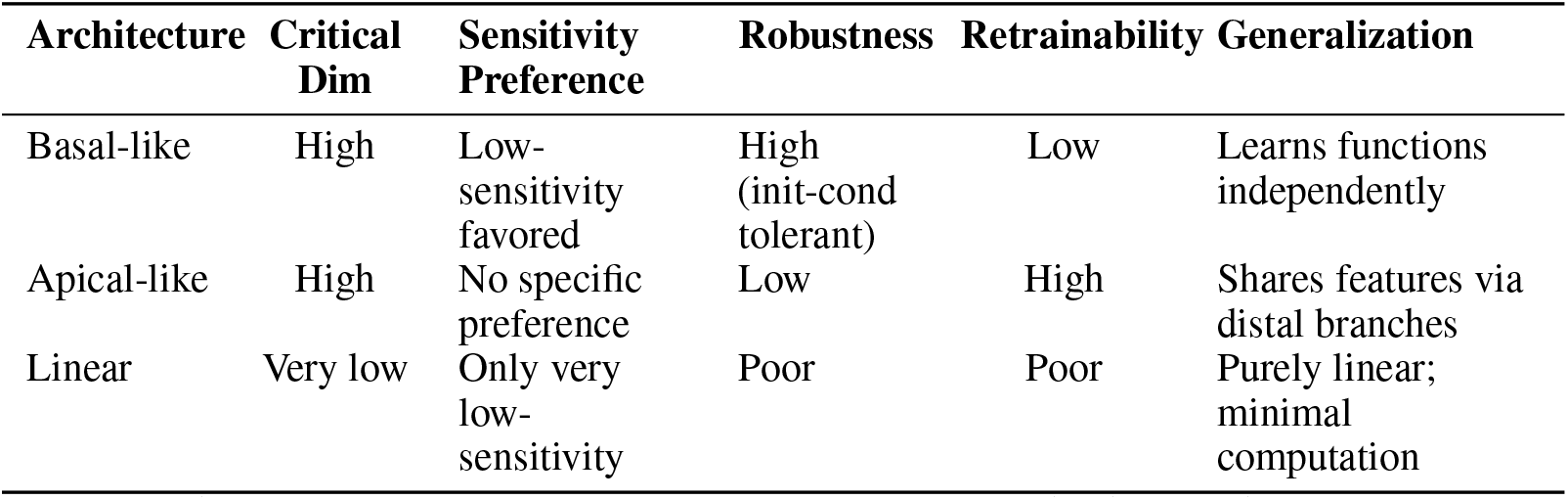
Summary of computational properties across dendritic architectures.

| Architecture | Critical Dim | Sensitivity Preference | Robustness | Retrainability | Generalization |
| --- | --- | --- | --- | --- | --- |
| Basal-like | High | Low-sensitivity favored | High (init-cond tolerant) | Low | Learns functions independently |
| Apical-like | High | No specific preference | Low | High | Shares features via distal branches |
| Linear | Very low | Only very low-sensitivity | Poor | Poor | Purely linear; minimal computation |

Beyond neuroscience, our findings offer new perspectives for artificial intelligence. Biological dendritic trees implement structured, resource-limited architectures that balance robustness and flexibility. This suggests engineered deep learning systems might benefit from constrained designs emulating dendritic specialization, especially in environments demanding continual adaptation without catastrophic forgetting. Understanding morphologys role in biological learning could inspire new strategies for more efficient, resilient machine learning models.

## References

Nathan W Gouwens, Staci A Sorensen, Jim Berg, Changkyu Lee, Tim Jarsky, Jonathan Ting, Susan M Sunkin, David Feng, Costas A Anastassiou, Eliza Barkan, et al. Classification of electrophysiological and morphological neuron types in the mouse visual cortex. Nature neuroscience, 22(7):1182–1195, 2019.

Nathan W Gouwens, Staci A Sorensen, Fahimeh Baftizadeh, Agata Budzillo, Brian R Lee, Tim Jarsky, Lauren Alfiler, Katherine Baker, Eliza Barkan, Kyla Berry, et al. Integrated morphoelectric and transcriptomic classification of cortical gabaergic cells. Cell, 183(4):935–953, 2020.

Nikolai C Dembrow, Scott Sawchuk, Rachel Dalley, Ximena Opitz-Araya, Mark Hudson, Cristina Radaelli, Lauren Alfiler, Sarah Walling-Bell, Darren Bertagnolli, Jeff Goldy, et al. Areal specializations in the morpho-electric and transcriptomic properties of primate layer 5 extratelencephalic projection neurons. Cell reports, 43(9), 2024.

Brian E Kalmbach, Rebecca D Hodge, Nikolas L Jorstad, Scott Owen, Rebecca de Frates, Anna Marie Yanny, Rachel Dalley, Matt Mallory, Lucas T Graybuck, Cristina Radaelli, et al. Signature morpho-electric, transcriptomic, and dendritic properties of human layer 5 neocortical pyramidal neurons. Neuron, 109(18):2914–2927, 2021.

Michael London and Michael Häusser. Dendritic computation. Annu. Rev. Neurosci., 28:503–532, 2005.

Romain Daniel Cazé, Mark Humphries, and Boris Gutkin. Passive dendrites enable single neurons to compute linearly non-separable functions. PLoS computational biology, 9(2):e1002867, 2013.

Brendan A Bicknell and Michael Häusser. A synaptic learning rule for exploiting nonlinear dendritic computation. Neuron, 109(24):4001–4017, 2021.

Albert Gidon, Timothy Adam Zolnik, Pawel Fidzinski, Felix Bolduan, Athanasia Papoutsi, Panayiota Poirazi, Martin Holtkamp, Imre Vida, and Matthew Evan Larkum. Dendritic action potentials and computation in human layer 2/3 cortical neurons. Science, 367(6473):83–87, 2020.

Chris Mingard, Joar Skalse, Guillermo Valle-Pérez, David Martínez-Rubio, Vladimir Mikulik, and Ard A Louis. Neural networks are a priori biased towards boolean functions with low entropy. arXiv preprint 1909.11522, 2019.

Claire Kenyon and Samuel Kutin. Sensitivity, block sensitivity, and -block sensitivity of boolean functions. Information and Computation, 189(1):43–53, 2004.

Michael Häusser and Bartlett Mel. Dendrites: bug or feature? Current opinion in neurobiology, 13 (3):372–383, 2003.

Alexandra Tzilivaki, George Kastellakis, and Panayiota Poirazi. Challenging the point neuron dogma: Fs basket cells as 2-stage nonlinear integrators. Nature communications, 10(1):3664, 2019.

Nelson Spruston. Pyramidal neurons: dendritic structure and synaptic integration. Nature Reviews Neuroscience, 9(3):206–221, 2008.

Richard M Karp. Reducibility among combinatorial problems. In 50 Years of Integer Programming 1958-2008: from the Early Years to the State-of-the-Art, pages 219–241. Springer, 2009.

Ingo Wegener. The complexity of Boolean functions. John Wiley & Sons, Inc., 1987.

Panayiota Poirazi and Bartlett W Mel. Impact of active dendrites and structural plasticity on the memory capacity of neural tissue. Neuron, 29(3):779–796, 2001.

Elizabeth Gardner and Bernard Derrida. Optimal storage properties of neural network models. Journal of Physics A: Mathematical and general, 21(1):271, 1988.

David Rubinstein. Sensitivity vs. block sensitivity of boolean functions. Combinatorica, 15(2): 297–299, 1995.

Elchanan Mossel, Ryan O’Donnell, and Krzysztof Oleszkiewicz. Noise stability of functions with low influences: invariance and optimality. In 46th Annual IEEE Symposium on Foundations of Computer Science (FOCS’05), pages 21–30. IEEE, 2005.

Gil Kalai. Boolean functions: influence, threshold and noise. In European Congress of Mathematics, volume 1, pages 85–110, 2016.

George Cybenko. Approximation by superpositions of a sigmoidal function. Mathematics of control, signals and systems, 2(4):303–314, 1989.

Donald R Jones, Cary D Perttunen, and Bruce E Stuckman. Lipschitzian optimization without the lipschitz constant. Journal of optimization Theory and Applications, 79:157–181, 1993.

Joerg M Gablonsky and Carl T Kelley. A locally-biased form of the direct algorithm. Journal of Global optimization, 21:27–37, 2001.

