## Supplemental File for "Bounds on the computational complexity of neurons due to dendritic morphology"

### Supplementary Material for: Bounds on the computational complexity of neurons due to dendritic morphology

**Anamika Agrawal***
Center for Data-Driven Discovery
Allen Institute
Seattle, WA 98109
Department of Neurobiology and Biophysics
University of Washington
Seattle, WA 98195


**Michael A. Buice**
Center for Data-Driven Discovery
Allen Institute
Seattle, WA 98109


#### 1 Supplemental Figures

*

Preprint.

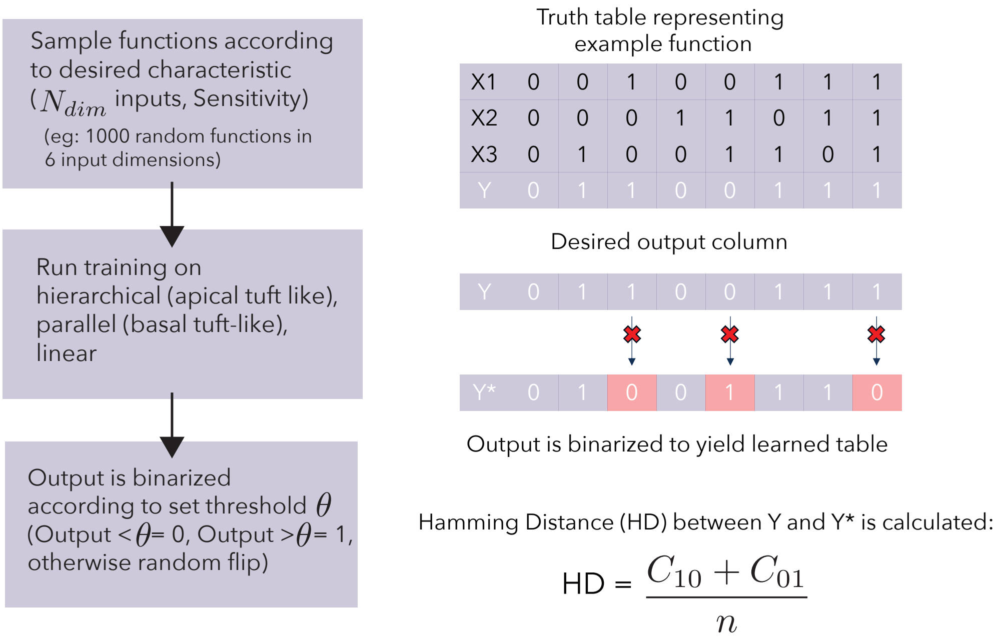


Figure 1: Schematic describing calculation of function learnability by an architecture through hamming distance of input truth table $Y_{in}$ and binarized output $Y_{out}$. The hamming distance (HD) quantifying the mismatch between the objective $Y$ and the learned output $Y*$ is defined as the sum of the number of 1-bits misclassified as 0 ($C_{10}$) and the number of 0-bits misclassified as 1 ($C_{01}$) divided by the total number of output bits.

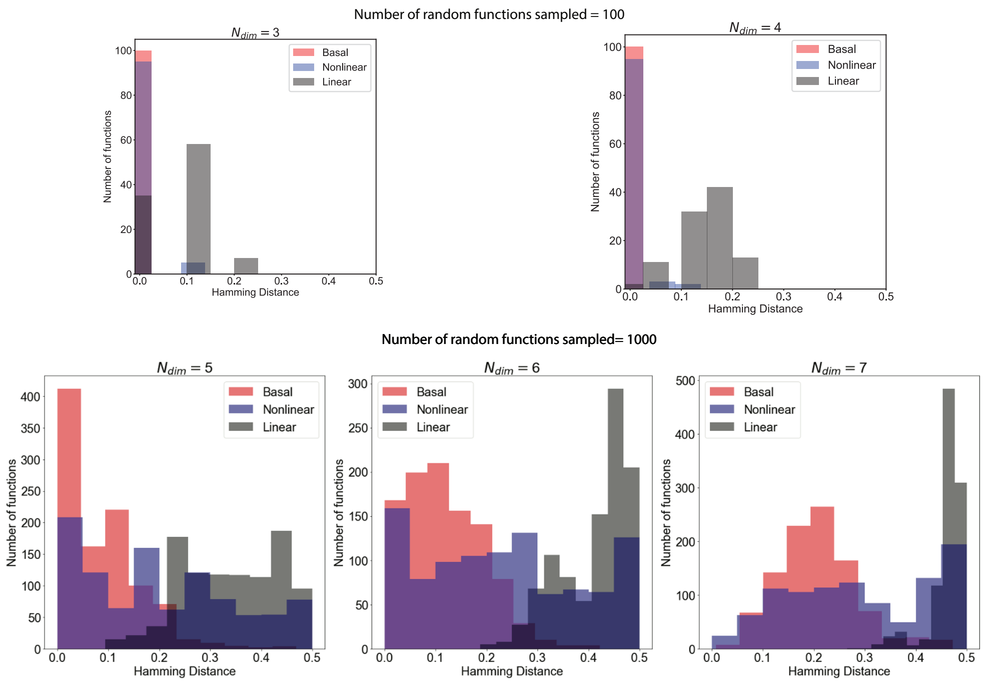


Figure 2: Performance Across input dimensions, highlights how linear neuron can't solve the typical functions. For $N_{dim} = 3, 4$; 100 random functions were sampled to be tested, for $N_{dim} = 5, 6, 7$, 1000 functions were sampled.

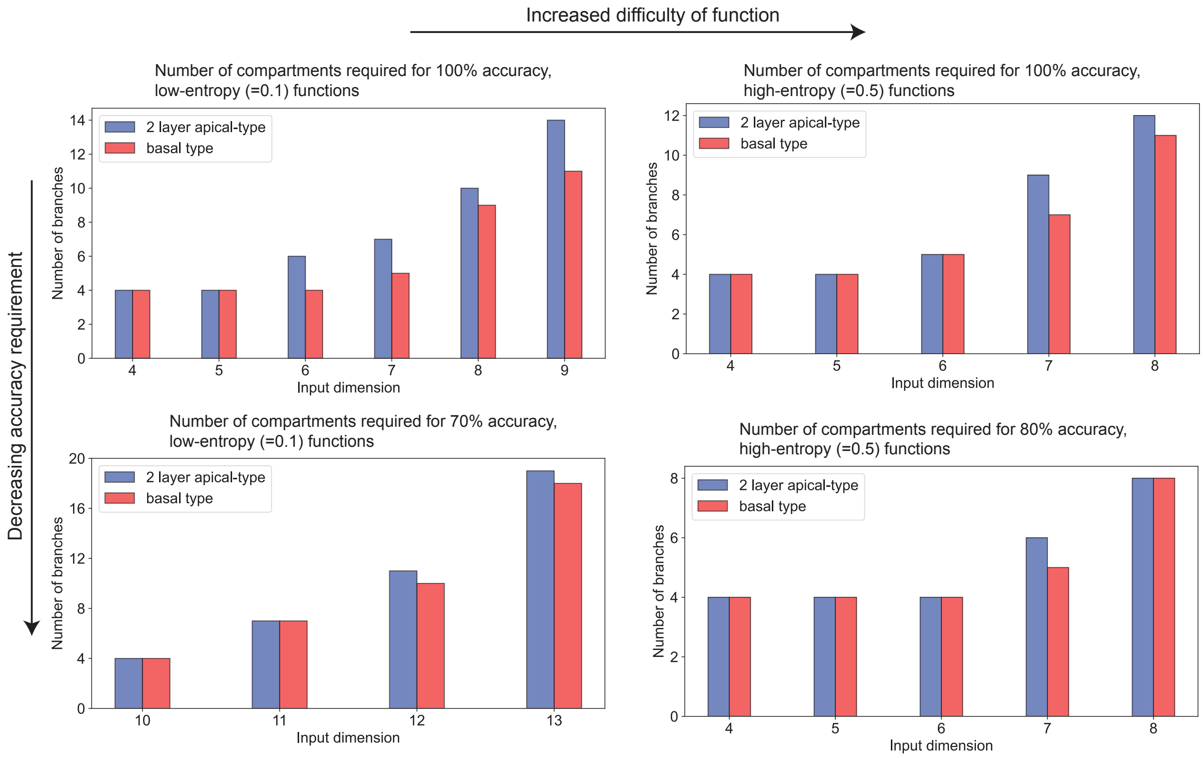


Figure 3: Compartment requirements scale exponentially. We tested the realizability of $n_{func} = 5$ randomly sampled functions in a given input dimension, with $n_{trials} = 5$ allowed to learn for each function. If an architecture is unable to learn any of the 5 functions in all of the trials, then the architecture is rejected and the compartment size is incremented by 1.There are 4 conditions tested: (i) low-entropy (low-difficulty) with high accuracy (ii) low-entropy with low accuracy (iii) high entropy, high accuracy (iv) high entropy, low accuracy

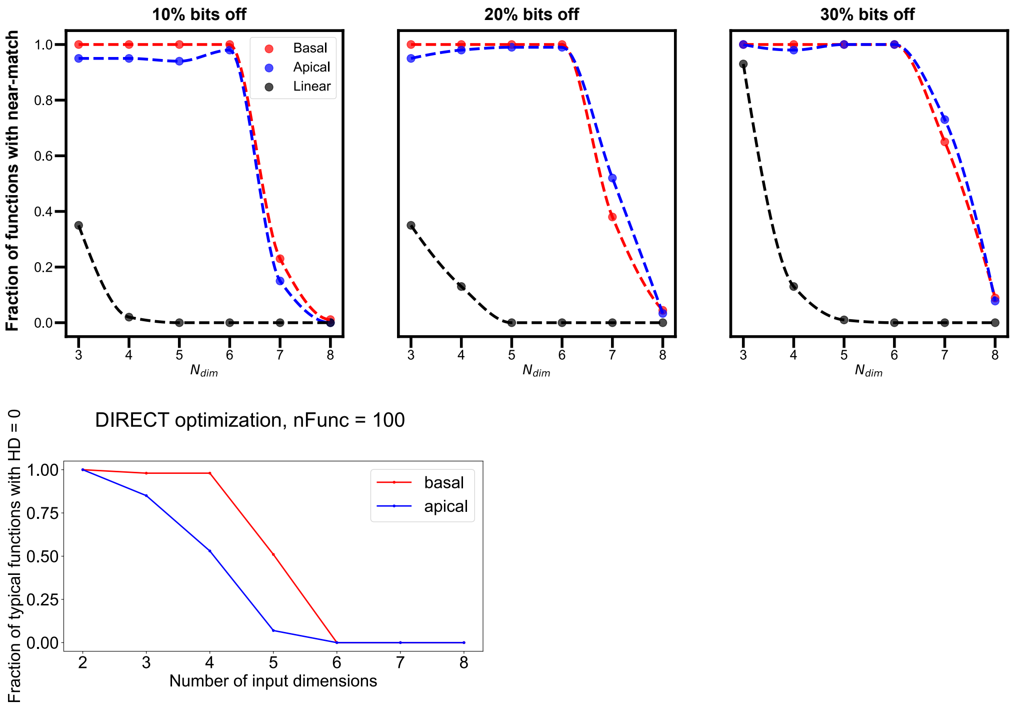


Figure 4: First row: The best case function realization fraction for each of the cell types, but with reduced requirements on accuracy. With reduced accuracy requirements. Second row: DIRECT based verification that there is a critical dimension beyond which functions become hard to realize, for both of the cell types.

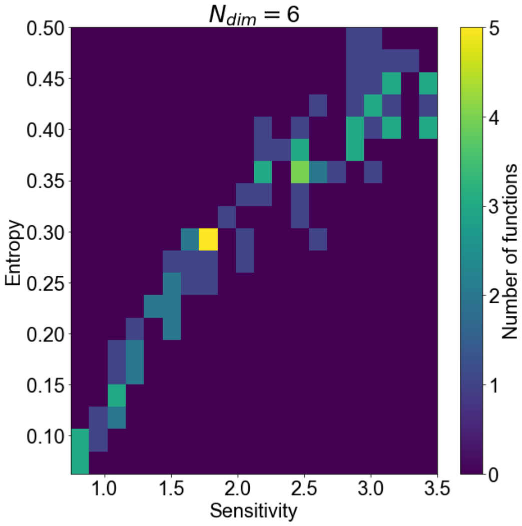


Figure 5: Entropy and Sensitivity have a monotonic relationship.

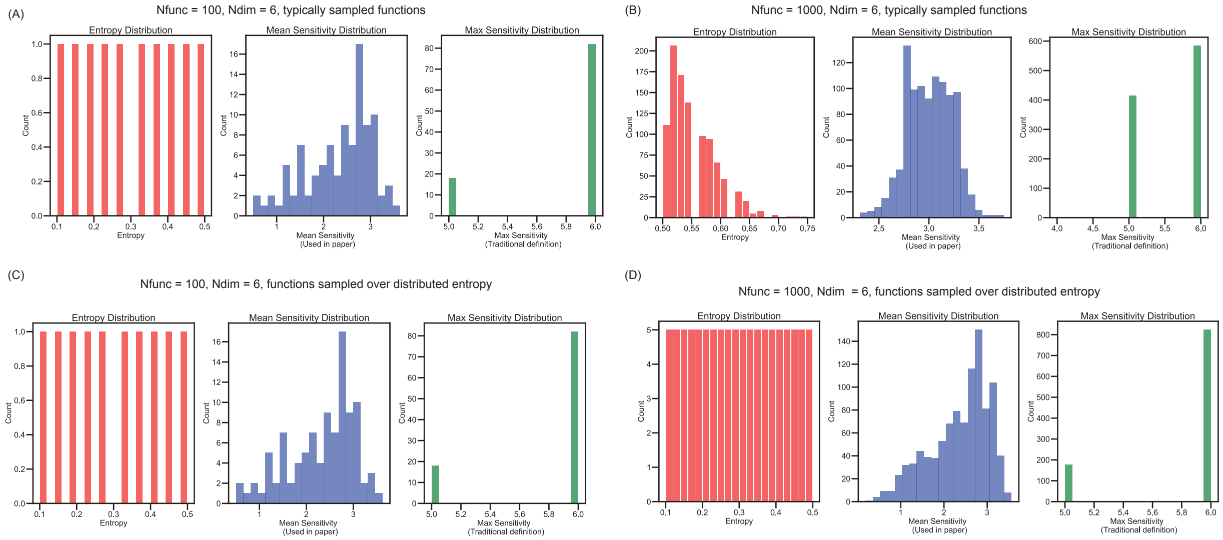


Figure 6: Figure demonstrates how the distribution of sensitivity changes from 100 typical functions to 1000 functions with distributed values of sensitivity, showing that the entropy-based sampling scheme successfully samples functions with a varying degree of sensitivity

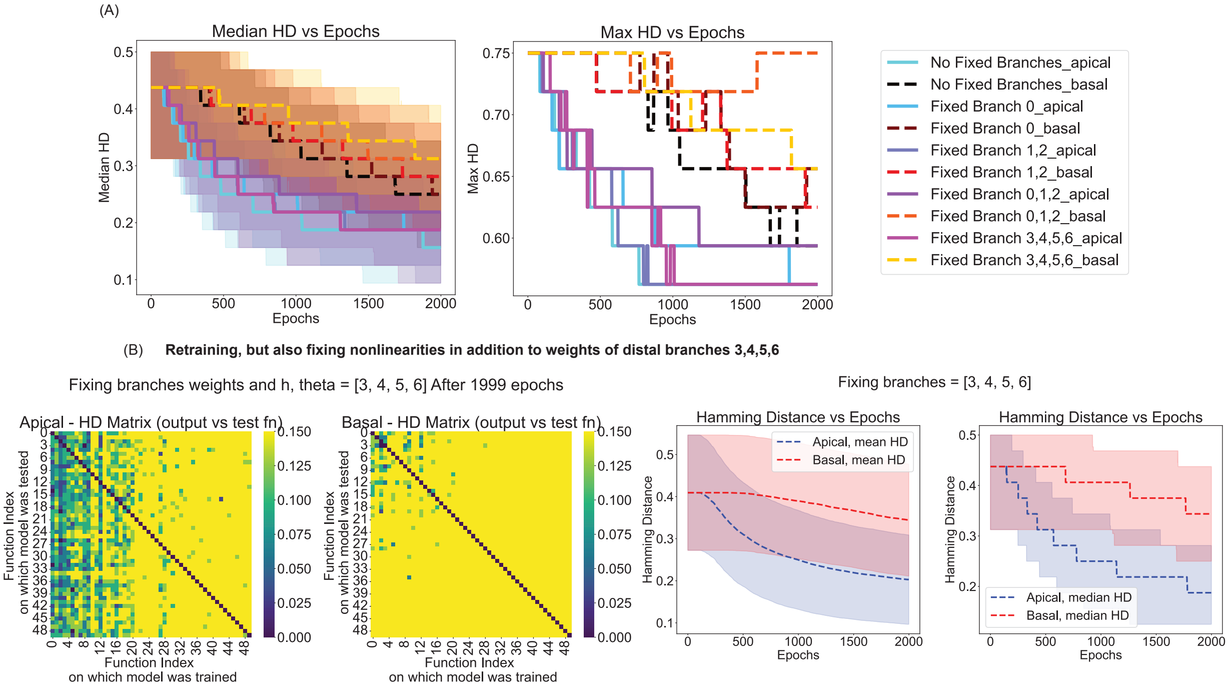


Figure 7: (A) Median and Max across $j$ for $HD_{j->i}(t)$ across $t$. (B) Flexible relearning is exhibited by apical type even when the non-linearities $h_b$ and $\theta_b$ are held fixed for branches $b = 3, 4, 5, 6$ (most distal branches/shallowest layer) during retraining.

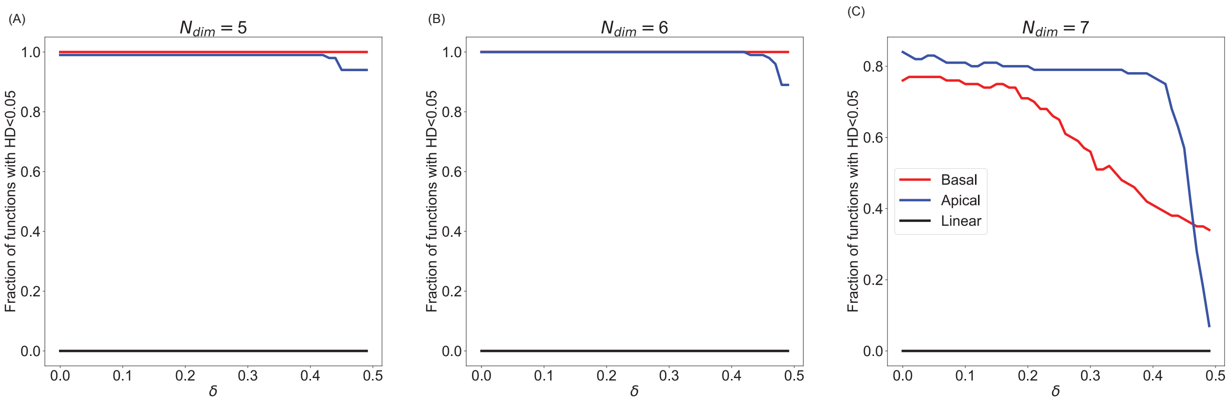


Figure 8: Changing $\delta$ (threshold for converting continuous output to binary output) changes the fraction of realized functions across dimensions. This threshold is referred to as $\theta$ in 1. At $N_{dim} = 7$ (the critical dimension for learning for both cells), the basal cell type is more sensitive to the threshold than the apical cell, highlighting two different methods of learning the boolean function mapping.

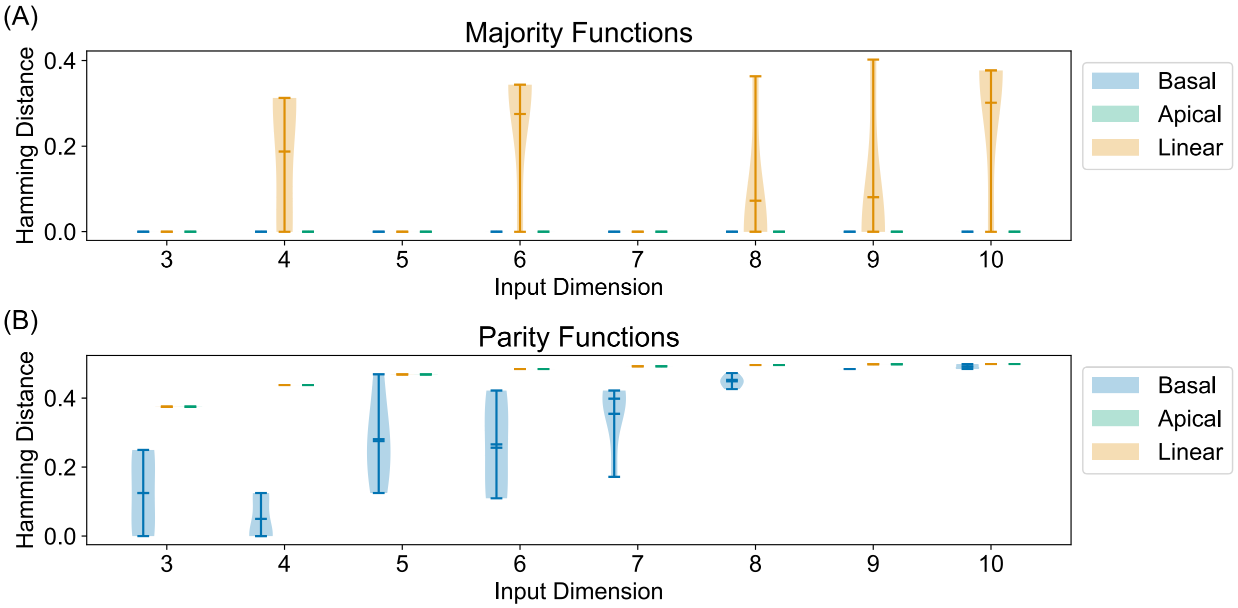


Figure 9: (A) Hamming distance while learning majority function (5 examples) in $N_{dim}$ (x axis) B) Hamming distance while learning parity function in $N_{dim}$ (x axis) Performance on (A) majority function is disproportionately good and (B) parity is disproportionately bad - highlighting inability to directly relate boolean circuit complexity classes to neural network circuit complexity.
